# How stimulation frequency and intensity impact on the long-lasting effects of coordinated reset stimulation

**DOI:** 10.1101/196188

**Authors:** Thanos Manos, Magteld Zeitler, Peter A. Tass

**Author notes:** (TM).

## Abstract

Several brain diseases are characterized by abnormally strong neuronal synchrony. Coordinated Reset (CR) stimulation was computationally designed to specifically counteract abnormal neuronal synchronization processes by desynchronization. In the presence of spike-timing-dependent plasticity (STDP) this may lead to a decrease of synaptic excitatory weights and ultimately to an anti-kindling, i.e. unlearning of abnormal synaptic connectivity and abnormal neuronal synchrony. The long-lasting desynchronizing impact of CR stimulation has been verified in pre-clinical and clinical proof of concept studies. However, as yet it is unclear how to optimally choose the CR stimulation frequency, i.e. the repetition rate at which the CR stimuli are delivered. This work presents the first computational study on the dependence of the acute and long-term outcome on the CR stimulation frequency in neuronal networks with STDP. For this purpose, CR stimulation was applied with Rapidly Varying Sequences (RVS) as well as with Slowly Varying Sequences (SVS) in a wide range of stimulation frequencies and intensities. Our findings demonstrate that acute desynchronization, achieved during stimulation, does not necessarily lead to long-term desynchronization after cessation of stimulation. By comparing the long-term effects of the two different CR protocols, the RVS CR stimulation turned out to be more robust against variations of the stimulation frequency. However, SVS CR stimulation can obtain stronger anti-kindling effects. We revealed specific parameter ranges that are favorable for long-term desynchronization. For instance, RVS CR stimulation at weak intensities and with stimulation frequencies in the range of the neuronal firing rates turned out to be effective and robust, in particular, if no closed loop adaptation of stimulation parameters is (technically) available. From a clinical standpoint, this may be relevant in the context of both invasive as well as non-invasive CR stimulation.

**Author Summary:** Abnormally strong neuronal synchronization is found in a number of brain disorders. To specifically counteract abnormal neuronal synchrony and, hence, related symptoms, Coordinated Reset (CR) stimulation was developed. CR stimulation employs basic plasticity and dynamic self-organization principles of the nervous system. Its fundamental goal is to induce long-lasting desynchronizing effects that persist cessation of stimulation. The latter are key to reducing side effects of invasive therapies such as deep brain stimulation. Furthermore, sustained stimulation effects pave the way for non-invasive neuromodulation treatments, where a few hours of stimulation delivered regularly or occasionally may provide substantial relief. Long-lasting CR-induced desynchronizing therapeutic effects have been verified in several pre-clinical and clinical studies. However, we here present the first computational study that systematically investigates the impact of key stimulation parameters on the stimulation outcome. Our results provide experimentally testable predictions that are relevant for pre-clinical and clinical studies. Furthermore, our results may contribute to stimulation techniques that enable to probe the functional role of brain rhythms in general.

## Introduction

Neuronal synchronization processes are relevant under normal as well as abnormal conditions [1]. A number of brain disorders are associated with abnormal neuronal synchrony, for example Parkinson’s disease [2–4], tinnitus [5–9] and epilepsy [10–12]. Neuronal dynamics and, in particular synchronization processes crucially depend on the patterns and types of neuronal connections [13–15]. For instance, according to computational studies it makes a significant difference whether neurons interact through gap-junctions or synapses [14, 15]. This is relevant for the emergence of different kinds of synchronization patterns [14–16] and epileptic seizures [17].

Connectivity and function are strongly connected and may undergo plastic changes throughout the life course [18]. The timing pattern of neuronal activity may strongly determine the strength of neuronal connections [19, 20]. Spike-timing-dependent plasticity (STDP) is a pivotal mechanism by which neurons adapt the strength of their synapses to the relative timing of their action potentials [21–25]. Based on seminal experimental studies [22–24] a series of computational studies focused on how adaptive coupling and activity dependent synaptic strength influence the collective neuronal dynamics [15, 17, 26–36]. In the presence of STDP a plethora of qualitatively different stable dynamical regimes emerge [15, 28, 36]. Qualitatively different stable dynamical states may actually coexist. In fact, multistability is a typical feature of neuronal networks and oscillator networks equipped with STDP. Multistability was found in different neural network models comprising different STDP models, e.g., in phase oscillator networks with both symmetric and asymmetric phase difference-dependent plasticity, a time continuous approximation of STDP [26, 28] as well as in phase oscillator networks with STDP [27] and in different types of neuronal networks with STDP [37–40] and other types of neural network models (e.g. [41–50] and references therein).

A number of computational studies were dedicated on desynchronizing synchronized ensembles or networks of oscillators or neurons [51–56]. The clinical need for stimulation techniques that cause desynchronization irrespective of the network’s initial state [57], thereby being reasonably robust against variations of system parameters and, hence, not requiring time-consuming calibration, motivated the design of Coordinated Reset (CR) stimulation [58, 59]. CR stimuli aim at disrupting in-phase synchronized neuronal populations by delivering phase resetting stimuli typically equidistantly in time, separated by time differences *T*_*s*_*N*_*s*_ where *T*_*s*_ is the duration of a *stimulation cycle*, and *N*_*s*_ is the number of active stimulation sites [58, 59]. The spatiotemporal sequence by which all stimulation sites are activated exactly once in a CR stimulation cycle is called the stimulation site sequence, or briefly *sequence*. Taking into account STDP [21–24] in oscillatory neural networks qualitatively changed the scope of the desynchronization approach: Computationally, it was shown that CR stimulation reduces the rate of coincident firing and, mediated by STDP, also decreases the average synaptic weight, ultimately preventing the network from generating abnormally increased synchrony [27]. This anti-kindling, i.e., unlearning of abnormally strong synaptic connectivity and of excessive neuronal synchrony, causes long-lasting sustained effects that persist cessation of stimulation [27, 37–39, 60, 61]. As shown computationally, anti-kindling can robustly be achieved in networks with plastic excitatory and inhibitory synapses, no matter whether CR stimulation is administered directly to the soma or through synapses [39, 62]. In line with these computational findings, long-lasting CR-induced desynchronization and/or therapeutic effects were accomplished with invasive as well as non-invasive stimulation modalities. Long-lasting desynchronization was induced by electrical CR stimulation in rat hippocampal slices rendered epileptic by magnesium withdrawal [63]. Electrical CR deep brain stimulation (DBS) caused long-lasting therapeutic after-effects in parkinsonian non-human primates [64, 65]. Bilateral therapeutic after-effects for at least 30 days were caused by unilateral CR stimulation delivered to the subthalamic nucleus (STN) of parkinsonian MPTP monkeys for only 2 h per day during 5 consecutive days [64]. In contrast, standard permanent high-frequency deep brain stimulation did not induce any sustained after-effects [64], see also [66]. In patients with Parkinson’s disease electrical CR-DBS delivered to the STN caused a significant and cumulative reduction of abnormal beta band oscillations along with a significant improvement of motor function [67]. Non-invasive, acoustic CR stimulation was developed for the treatment of patients suffering from chronic subjective tinnitus [62, 68]. In a proof of concept-study acoustic CR stimulation caused a statistically and clinically significant sustained reduction of tinnitus symptoms [68–70] together with a concomitant decrease of abnormal neuronal synchrony [68, 71], abnormal effective connectivity [72] as well as abnormal cross-frequency coupling [73] within a tinnitus-related network of brain areas.

So far, the pre-clinical [68, 74] and clinical [64, 67] proof of concept studies for invasive and non-invasive CR stimulation were driven by computationally derived hypotheses and predictions. Theoretically predicted phenomena and mechanisms, such as long-lasting stimulation effects [27, 37, 39, 60], cumulative stimulation effects [61], and improvement by weak stimulus intensity [75] were verified based on dedicated theory-driven study protocols for pre-clinical and clinical proof of concepts [64, 67, 68, 74].

We here set out to investigate the impact of the CR stimulation frequency and intensity on the effects during stimulus delivery (so-called *acute effects*), on transient effects emerging directly after cessation of stimulation (so-called *acute after-effects*), and on effects outlasting cessation of stimulation (so-called *sustained after-effects*). The ultimate goal of this study is to improve the calibration of CR stimulation, in particular, by providing computationally generated predictions that can be tested in subsequent pre-clinical and clinical studies. The computational study presented here is organized around three hypotheses:

*Hypothesis #1: Due to the inherently periodic structure of CR stimulation the relation between CR stimulation frequency and the spontaneous neuronal firing rates (prior to stimulation) matters.* Periodic delivery of CR stimuli with fixed sequence basically constitutes a time-shifted entrainment of the different neuronal subpopulations [58, 59]. A particular closed loop embodiment of CR stimulation, periodic stimulation with demand-controlled length of high-frequency pulse train, is basically a time-shifted entrainment of the different neuronal subpopulations with stimulus intensities adapted to the amount of undesired synchrony [58, 59]. Accordingly, the duration of a stimulation cycle was selected to be reasonably close to the mean period of the synchronized neuronal oscillation [58, 59]. In STDP-free networks of Kuramoto [76] and FitzHugh-Nagumo [77] model neurons the impact of CR stimulation intensity and frequency on the desynchronizing outcome of CR was studied in detail [75].

*Hpothesis #2: Different embodiments of CR stimulation may differ with respect to effect size and robustness.* In a series of computational studies [39, 40, 58, 59, 75, 78, 79] and in all pre-clinical [63, 64] and clinical studies [67–72] performed so far, CR was applied either with fixed sequences or *rapidly varying sequences (RVS)*, where the sequence was randomly varied from cycle to cycle. In a recent computational study, it was shown that at intermediate stimulation intensities the CR-induced anti-kindling effect may significantly be improved by CR with *slowly varying sequences (SVS)*, i.e. by appropriate repetition of the sequence with occasional random switching to the next sequence [78]. However, this study was not performed for a larger range of CR stimulation frequencies. By definition, SVS CR stimulation features significantly more periodicity of the stimulus pattern. Accordingly, the dependence of resonance and/or anti-resonance effects on the CR stimulation frequency might be more pronounced for SVS CR as opposed to RVS CR.

*Hypothesis #3: Pronounced acute effects might provide a necessary, but not sufficient condition for pronounced sustained after-effects.* In a pre-clinical study in Parkinsonian monkeys with CR-DBS delivered at an optimal and a less favorable intensity, it was shown that long and pronounced acute therapeutic after-effects coincide with long-lasting, sustained after-effects [68]. However, according to computational studies the relationship between acute after-effects and sustained long-lasting effects might be more involved, at least for particular parameter combinations [78].

Related to these hypotheses, to assess the robustness of CR stimulation against initial network conditions we performed our numerical simulations for different network initializations, respectively. In this study we did not systematically vary the stimulation duration. Rather, based on a pre-series of numerical simulations, we here used a fixed stimulation duration that is reasonably short, but nevertheless enabled to robustly achieve an anti-kindling for properly selected values of stimulation frequency and intensity. In fact, our goal was to find stimulation parameters enabling short, but notwithstanding effective CR stimulation. Keeping the stimulation duration at moderate levels may be beneficial for applying the CR approach to different invasive as well as non-invasive stimulation modalities. For instance, standard DBS, i.e. permanent electrical high-frequency pulse train stimulation delivered to dedicated target areas through implanted depth electrodes, used for the treatment of, e.g., Parkinson’s disease [80–82] may cause side effects. If side effects are caused by stimulation of non-target tissue, they may be reduced by adapting the spatial extent of the current spread to the target’s anatomical borders by appropriate electrode designs as introduced, e.g., by [83–85], in particular, to spatially tailor stimuli by means of directional DBS [86–91]. However, some side effects may at least partly be caused by stimulating the target region itself [92, 93]. Accordingly, no matter how precisely stimuli are delivered to DBS targets, the amount of stimulation should be decreased as much as possible. Another example refers to non-invasive applications of CR. In general, non-invasive CR stimulation requires the patients’ compliance to actually pursue treatment prescriptions. Obviously, patients might prefer shorter treatment sessions.

To come up with favorable combinations of stimulation parameters, in our numerical analysis we used different data analysis methods, e.g. macroscopic measures assessing the average amount of the population’s synchrony and synaptic connectivity. These measures are appropriate to demonstrate relevant stimulation effects, such as stimulation-induced transitions from pronounced neuronal synchrony to desynchronized states.

In summary, in this paper we first explain the computational model and analysis methods. We then apply RVS CR stimulation in a wide parameter range of stimulation frequencies and intensities. We repeat the same analysis for SVS CR stimulation and investigate the differential characteristics of RVS CR and SVS CR with respect to efficacy and robustness. Finally, we analyze the relationship between stimulation-induced acute effects and after-effects. Our results provide the foundation for the development of novel control techniques that will be the topic of a forthcoming study.

## Materials and Methods

### The Hodgkin-Huxley Spiking Neuron Model

In this study we use the conductance-based Hodgkin-Huxley neuron model [94] for the description of an ensemble of spiking neurons. The set of equations and parameters are (see also [95, 96]):

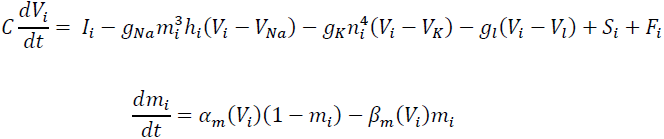

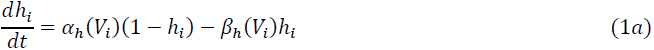

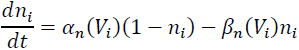

The variable *V*_*i*_, with *i* = 1, …, *N*, describes the membrane potential of neuron *i*, and:

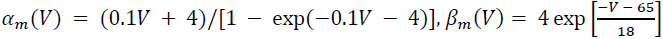

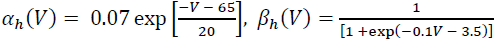

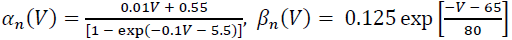

The total number of neurons is *N* = 200, while *g*_*Na*_ = 120 mS/cm^2^, *g*_*K*_ = 36 mS/cm^2^, *g*_*l*_ =0.3 mS/cm^2^ are the maximum conductance per unit area for the sodium, potassium and leak currents respectively. The constants *V*_*Na*_ = 50 mV, *V*_*K*_ = −77 mV and *V*_*l*_ = −54.4 mV refer to the sodium, potassium and leak reversal potentials respectively. *C* is the constant membrane capacitance (*C* = 1 µF/cm^2^), *I*_*i*_ the constant depolarizing current injected into neuron *i*, determining the intrinsic firing rate of the uncoupled neurons. For the realization of different initial networks, we used random initial conditions drawn from uniform distributions, i.e. *I*_*i*_∈ [*I*_0_-σ_*I*_, *I*_0_+ σ_*l*_] (*I*_0_ = 11.0 µS/cm^2^ and σ_*l*_ = 0.45 µS/cm^2^), *h*_*i*_, *m*_*i*_, *n*_*i*_∈ [0, 1] and *V*_*i*_∈ [-65, 5] mV. The initial values of the neural synaptic weights *c*_*ij*_ are picked from a normal distribution *N*(µ_*c*_ = 0.5 µA/cm^2^, σ_*c*_ = 0.01 µA/cm^2^) (see also [39, 78] for more details). Hence, in this setup the neurons are not identical. The *S*_*i*_ term refers to the internal synaptic input of the neurons within the network to neuron *i*, while *F*_*i*_ represents the current induced in neuron *i* by the external CR stimulation.

### Network and Neuron Coupling Description

The *N* = 200 spiking Hodgkin-Huxley neurons are placed on a ring and the *N*_*s*_ = 4 stimulations sites are equidistantly placed in space at the positions of neurons *i* = 25, 75, 125, 175. The neurons interact via excitatory and inhibitory chemical synapses by means of the postsynaptic potential (PSP) *s*_*i*_ which is triggered by a spike of neuron *i* [21, 97] and modelled using an additional equation (see also [98, 99]):

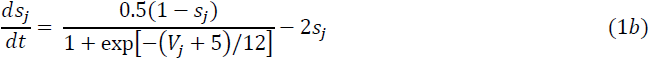

Initially we draw *s*_*i*_∈ [0, 1] (randomly from a uniform distribution) and then, the coupling term S_i_ from Eq (1a) (see [39]) contains a weighted ensemble average of all postsynaptic currents received by neuron *i* from the other neurons in the network and is described by the following term:

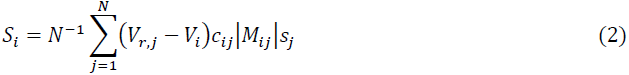

where *V*_*r*,*j*_ is the reversal potential of the synaptic coupling (20 mV for excitatory and –40 mV for inhibitory coupling) and *c*_*ij*_ is the synaptic coupling strength from neuron *j* to neuron *i*. There are no neuronal self-connections within the network (*c*_*ii*_ = 0 mS/cm^2^). The term *M*_*ij*_, which describes the spatial profile of coupling between neurons *i* and *j*, is given by:

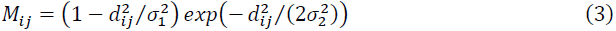

It has the form of a Mexican hat [100–102] and defines the strength and type of neuronal interaction: strong short-range excitatory (*M*_*ij*_ > 0) and weak long-range inhibitory interactions (*M*_*ij*_ < 0). Here *d*_*ij*_ = *d*|*i* - *j*| is the distance between neurons *i* and *j*, while

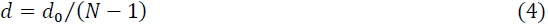

determines the distance on the lattice between two neighboring neurons within the ensemble, *d*_0_ is the length of the neuronal chain (*d*_0_ = 10), σ _1_ = 3.5, and σ _2_ = 2.0. In order to limit boundary effects, we consider that the neurons are distributed in such a way that the distance *d*_*ij*_ is taken as: *d* · *min*(|*i* - *j*|, *N* - |*i* - *j*|) when the *i*, *j* > *N*/2.

### Spike-Timing-Dependent Plasticity

We follow the concepts described in [22, 23], regarding the synaptic coupling strengths change dependence on the precise timing of pre- and post-synaptic spikes. Hence, we consider all the synaptic weights *c*_*ij*_ to be dynamic variables that depend on the time difference (∆*t*_*ij*_) between the onset of the spikes of the post-synaptic neuron *i* and pre-synaptic neuron *j* (denoted by *t*_*i*_ and *t*_*j*_ respectively). Then the STDP rule for the change of synaptic weights is given by [23, 39]:

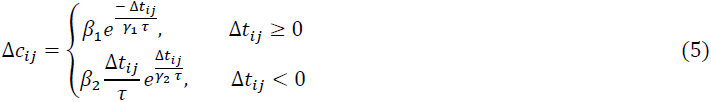

Whereβ_1_ = 1, β_2_ = 16, γ_1_ = 0.12, γ_2_ = 0.15, τ ms and δ = 0.002. According to the value of ∆*t*_*ij*_, the synaptic weight *c*_*ij*_ is updated in an event-like manner, i.e. we add or subtract an increment δ · ∆*c*_*ij*_ for excitatory or inhibitory connections respectively, with learning rate δ > 0 every time a neuron spikes. Furthermore, we restrict the values of *c*_*ij*_ on the interval [0,1] mS/cm^2^ for both excitatory and inhibitory synapses, ensuring in this way that their strengthening or weakening remains bounded. The maximal inhibitory synaptic weight *cm*𝑎*x* was set to be 1 in all our stimulations. However, a more detailed investigation about the effect and variation of this value was performed in [78] where when increasing *cm*𝑎*x* of the inhibitory neurons no significant impact was observed regarding (de)synchronization effects accompanied with a lower average network connectivity.

### Coordinated Reset Stimulation

Coordinated Reset (CR) stimulation was applied to the neuronal ensemble of *N* spiking Hodgkin-Huxley neurons. This was done sequentially via *N*_*s*_(=4 in this study) equidistantly spaced stimulation sites [58]: one stimulation site was active during *T*_*s*_/*N*_*s*_, while the other stimulation sites were inactive during that period. After that another stimulation site was active during the next *T*_*s*_/*N*_*s*_ period. All *N*_*s*_ stimulation sites were stimulated exactly once within one stimulation ON-cycle. Therefore, the duration of each ON-cycle is *T*_*s*_(in ms). The spatiotemporal activation of stimulation sites is represented by the indicator functions *ρ*_*k*_(*t*) (k∊ {1, …, *N*}):

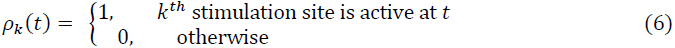

The stimulation signals induced single brief excitatory post-synaptic currents. The evoked time-dependent normalized conductances of the postsynaptic membranes are represented by α-functions given in [96]:

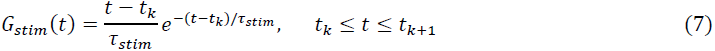

Here τ_*stim*_ = *T*_*s*_/(6*N*_*s*_) denotes the time-to-peak of *G*_*stim*_ and *t*_*k*_ is the onset of the *kth* activation of the stimulation site. Note that the period (or frequency) through the τ_*stim*_ parameter of the CR stimulation determines not only the onset random timing of each single signal but also its temporal duration. The spatial spread of the induced excitatory postsynaptic currents in the network is defined by a quadratic spatial decay profile (see [96] for more details) given as a function of the difference in index of neuron *i* and the index *x*_*k*_ of the neuron at stimulation site *k*

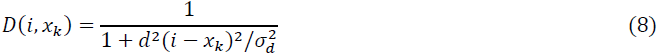

with *d* the lattice distance between two neighboring neurons as defined in Eq (4) and σ_*d*_=0.08 the spatial decay rate of the stimulation current. Thus the total stimulation current from Eq (1) is expressed by the following equations:

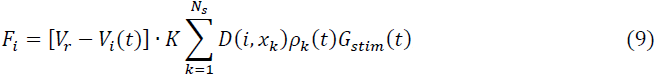

where *V*_*r*_ = 20 mV denotes the excitatory reverse potential, *V*_*i*_ the membrane potential of neuron *i*, *K* the stimulation intensity, and *ρ*, *G, D* are given by Eqs (6), (7), and (8) respectively. For the RVS CR stimulation, sequences are randomly chosen from a set of *N*_*s*_!(=24) different possible sequences during each ON-cycle (an example is shown in Fig 1A for CR stimulation period *T*_*s*_ = 10 ms for the first 90 ms of an activated CR period). Each newly drawn sequence is indicated by a different color and lasts exactly one ON cycle. On the other hand, for the SVS-*l* CR stimulation, one first randomly picks a sequence and repeats it *l* times before switching to another one, as shown by the example in Fig 1B (again for *T*_*s*_ = 10 ms) for *l* = 4. The administered stimulation protocol consists of *m*: *n* = 3: 2 CR ON-OFF cycles (see [39, 75, 78]). Depending on the *T*_*s*_ value, more (or less) ON-cycles may be administered within a fixed time interval. In this panel, the total time spans up to two completed ON-and OFF cycles (up to ∼125 ms in this case) and the color changes at each new sequence.

**Fig 1.**
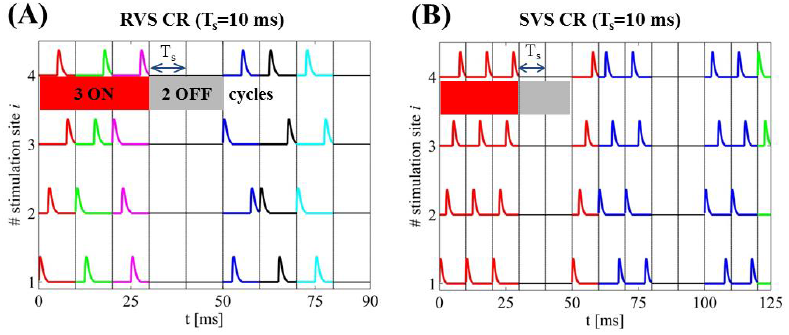
Time evolution of CR stimulation signals. (A) RVS CR stimulation signal with period *T*_*s*_ = 10 ms for the first 90 ms of an activated CR period. The vertical lines indicate the successive ON- and OFF cycles and the temporal distance between two successive vertical lines correspond to the period *T*_*s*_ of each cycle (every stimulation site is activated exactly once during the ON cycles). A change of color indicates a change of sequence. (B) SVS-4 CR stimulation signal with the same period but here the total time spans up to two completed ON-and OFF cycles ∼125ms) while the color changes as a new sequence is drawn.

### Macroscopic measurements and statistical tools

The synaptic weights, being affected by the STDP and the different intrinsic periods of the neurons, change dynamically in time. One efficient way to measure the strength of the coupling within the neuronal population at time *t* is given by the following synaptic weight (averaged over the neuron population):

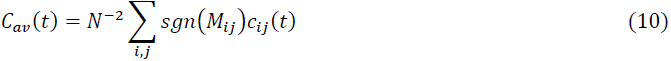

where *M*_*ij*_ is defined in Eq (3) and *sgn* is the sign-function. Furthermore, one may additionally measure the degree of the neuronal synchronization within the ensemble, using the order parameter [76, 103]:

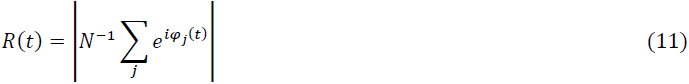

Where φ_j_(t) = 2π(t - *t*_*j*,*m*_)/ (*t*_*j*,*m*+1_- *t*_*j*,*m*_) for *t*_*j*,*m*_≤ *t* < *t*_*j*,*m*+1_ is a linear approximation of the phase of neuron *j* between its *mth* and (*m* + 1)*th* spikes at spiking times *t*_*j*,*m*_ and *t*_*j*,*m*+1_. *R*(*t*) is influenced by the synaptic weights, as the latter are time dependent due to the STDP. The order parameter *R* measures the extent of phase synchronization in the neuronal ensemble and takes values between 0 (absence of in-phase synchronization) and 1 (perfect in-phase synchronization).

In our numerical calculations, we estimate *C*_*av*_[see Eq (10)] and *R*_*av*_. The latter quantity is averaged over the last 100 · *T*_*s*_. Whenever we plot the order parameter versus time, we determine the moving average < *R* > over a time window of 400 · *T*_*s*_, because of the presence of strong fluctuations. For the statistical description and analysis of the non-Gaussian distributed *C*_*av*_ and *R*_*av*_ data (*n* = 11 samples), we use the median as well as the Inter-Quartile Range (IQR) [104]. The IQR measures the statistical dispersion, namely the width of the middle 50% of the distribution and is represented by the box in a boxplot. It is also used to determine outliers in the data: an outlier falls more than 1.5 times IQR below the 25% quartile or more than 1.5 times IQR above the 75% quartile.

### Simulation Description

In this study, for each initial network of *N* = 200 non-identical-neurons and parameter conditions (or simply “network”), we apply RVS and SVS CR signals (different per network). For each network, the initial conditions for each neuron were randomly drawn from random distributions as given in the *Hodgkin-Huxley Spiking Neuron Model* subsection. We start the simulation with an equilibration phase, which lasts 2 s. Later on, we evolve the network under the influence of STDP (which will be present until the end of the simulation). We then integrate the network for 60 s with STDP without any external stimulation yet, where a rewiring of the connections takes place, resulting in a strongly synchronized state. Next, we apply CR stimulation for 128 s (resetting the starting time to *t* = 0 s). During this CR-on period three stimulation ON-cycles (the stimulation is on) alternated with two OFF-cycles (the stimulation is off) as in the example stimulation signals shown in Fig 2. Each ON- and OFF-cycle lasts *T*_*s*_. After 128 s the CR stimulation ceases permanently and we continue the evolution of the CR-off period for extra 128 s.

**Fig 2.**
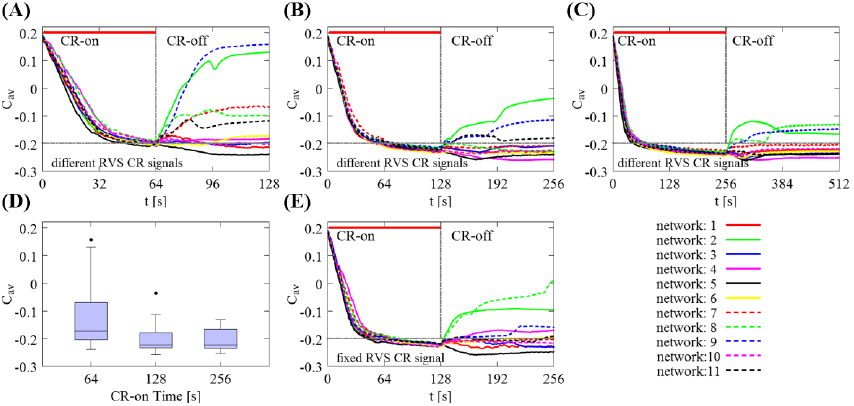
Impact of the total CR-on time on the mean synaptic weight *C*_*av*_ for different initial random networks and RVS CR. (A) Time evolution of the *C*_*av*_ for different total CR-on time durations, *t* = 64 s, (B) *t* = 128 s (this is the standard CR-on period used throughout the paper) and (C) *t* = 256 s. In all these cases, 11 different initial networks were stimulated with different RVS CR stimulation signals during the CR-on period. The thick red horizontal lines indicate the CR-on/off stimulation periods (the end is marked with a vertical gray line) while the horizontal gray dashed lines are visual cues for mutual comparison. (D) Boxplots of the mean synaptic weights presented in (A)-(C), showing the median values (black lines within the boxes). The box frames depict the middle 50%, the upper and lower whiskers the first and last 25% respectively while the outliers (black dots) are set as 1.5 times the length of the box (above/below). There is no statistically significant difference between the data sets at t = 128 s and t = 256 s (*p* = 0.8955 two-sided Wilcoxon rank sum test). The total CR-on/off time is twice as long as the CR-on period. (E) An identical RVS CR stimulation signal (the one of network 1) was used for all 11 initial networks for *t* = 128 s [comparison with (B)]. In all cases, the CR stimulation intensity is *K* = 0.20 with period *T*_*s*_ = 10 ms.

In order to probe and chart the CR stimulation intensity and frequency parameter space, we restrict the CR stimulation intensity to values in the interval (*K*∈ [0.20, …, 0.50]). This particular choice is based on our previous experience and numerical studies (see e.g. [39, 78]) where it was found that weaker intensities were not able to sufficiently desynchronize the neuron ensemble while larger intensities did not significantly improve (or sometimes even worsen) the outcome of RVS and SVS CR stimulation signals. We then set an initial-central value for the CR stimulation period (that defines the initial/starting frequency) which in principle is selected close to the intrinsic firing rate of the strongly synchronized network. In this case, and before applying the CR stimulation, the intrinsic firing rate of the network is ∼71 Hz which corresponds to *T*_*s*_ ≈ 14 ms. Hence, we begin with the CR stimulation period *T*_0_ = 16 ms which gives an initial stimulation frequency f_0_ = 1/*T*_0_(in a similar manner just like in [39, 78] and adjusted to a value close to the intrinsic one). Then we define such a period interval [*T*_*s*_*min*, *T*_*s*_*m*𝑎*x*] in ms (*T*_*s*_: integer) that allows us to create an “approximately” equidistant grid on the frequency space: f_stim_∈[25%f_0_, …, 175%f_0_]. This initial *T*_0_- value is also well studied for different types of signal patterns aiming to optimize the CR effect with the use of different type of CR stimulation sequences (see e.g. [78]). Then, we define the ratio (%) of CR sequence frequency per ON-cycle (f_stim_) over the frequency of the reference stimulation frequency (f_0_ = 62.5 Hz, *T*_0_ = 16 ms) as *r*_0_ = (f_stim_/f_0_) · 100 and we end up in studying the intensity and frequency-ratio (*K*, *r*_0_) - parameter space. In Table 1, we show the conversion between the stimulation frequency-ratio and period. For comparison reasons, we also give the corresponding ratios *r*_int_(%) of CR stimulation frequency per ON-cycle (f_stim_) over the frequency of the *intrinsic firing rate* of the network frequency (f_int_ = 71.4 Hz, *T*_int_=14 ms) without any external stimulation.

**Table 1.**
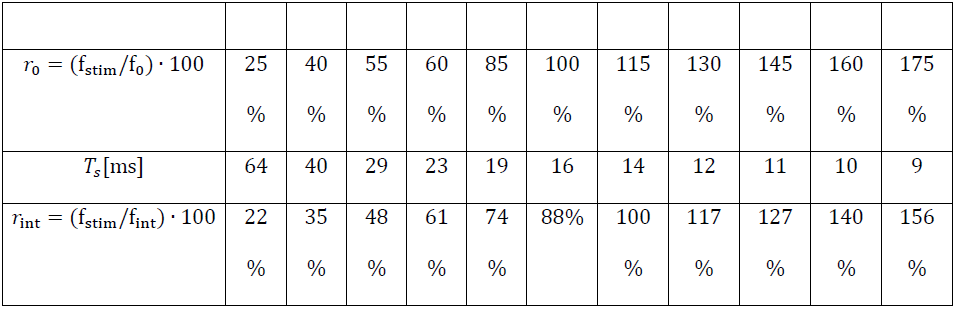
Conversion between the stimulation frequency and period.

In the first row, we show the ratio *r*_0_(%) of the CR sequence frequency per ON-cycle (f_stim_) over the frequency of the reference stimulation frequency (f_0_ = 62.5 Hz, *T*_0_ = 16 ms) which is used for providing f_stim_-values which are distributed in an “approximately” equidistant grid on the frequency space: f_stim;_∈ [25%f_0_, …, 175%f_0_]. Based on these values, we define the period *T*_s_(second row) of the CR sequences. These are the two descriptions used broadly throughout the paper. In the last row, we show additionally - and only for comparison reasons - the corresponding ratios *r*_int_(%) of CR sequence frequency per ON-cycle (f_stim_) over the frequency of the *intrinsic firing rate* of the network frequency (f_int_ ≈ 71.4 Hz, *T*_int_ ≈ 14ms) without any external stimulation.

## Results

### Impact of CR Stimulation Duration and Signals on Different Initial Networks

Before presenting the core of our findings, let us first start by discussing how the RVS CR stimulation duration affects the long-lasting anti-kindling of different initial randomly chosen networks. In Fig 2, we show the evolution of the mean synaptic weight *C*_*av*_ as a function of time for different total CR-on time durations: *t* = 64 s (Fig 2A), *t* = 128 s (Fig 2B), and *t* = 256 s (Fig 2C). 128 s is the standard CR-on period used throughout the paper. The CR stimulation intensity is *K* = 0.20, and the period *T*_*s*_ = 10 ms. A general trend appears in this sequence of panels, i.e. the longer the CR stimulation lasts, less spread of the *C*_*av*_ regarding the long-lasting anti-kindling effect is observed after stimulation offset. This is shown in Fig 2D with boxplots. The last box (corresponding to *t* = 256 s of total CR-on period) has no outliers and shows a more “uniform” long-lasting effect (as shown in Fig 2C) for all 11 network initializations, not only during the CR-on period but also afterwards during the CR-off period. However, there is no statistically significant decrease of the median of the *C*_*av*_ from *t* = 64 s to *t* = 128 s (right-sided Wilcoxon rank sum test [105], *p* = 0.0209, 5% significance level). Moreover, the median value of the *C*_*av*_ does not change significantly between *t* = 128 s (Fig 2B) and *t* = 256 s (Fig 2C, both-sided Wilcoxon rank sum test, *p* = 0.8955). Hence, the intermediate stimulation duration *t*= 128 s provides fairly good results. Furthermore, for considerably larger stimulation durations the anti-kindling is typically, but not always more pronounced. From a clinical standpoint, it is desirable to achieve reasonably pronounced stimulation effects without excessive stimulation durations. Accordingly, in this computational study we choose *t* = 128 s as total CR-on time, and *t* = 256 s as total CR-on/off time.

For the different simulations, we use different random initial networks and CR signals. For the sake of generality, we do not pick any optimal combination of random initial network and RVS CR stimulation signal that would induce a favorable or biased behavior. This is to assess whether CR effects are robust with respect to different initial conditions.

Fig 2B shows a typical example where 11 different random stimulation signals where applied to 11 different initial networks during the CR-on period, with CR stimulation intensity *K* = 0.20 and stimulation period *T*_*s*_ = 10 ms. The CR-on/off period lasts 128 ms respectively. During the CR-on period the mean synaptic weights *C*_*av*_ evolve in a similar manner for all networks, with little deviations between the different curves. They reach approximately the same small value at the end of the CR-on period. The latter corresponds to weak excitatory synaptic connectivity and, in most cases in this paper, to globally well-desynchronized states. However, the post-stimulation dynamics of *C*_*av*_ may be quite diverse. Some networks retain their weak average connectivity while others, like network 2 and 9 (Fig 2B) relapse back to states with strong synaptic connectivity. Next, we study what happens if we fix the CR stimulation signal for the 11 different initial networks (Fig 2E). The results are similar to Fig 2B: The outcome at the end of the CR-on period is fairly uniform, while the post-stimulation dynamics of *C*_*av*_ is diverse. Replacing one random external stimulation signal by another one may improve the long-term outcome in some cases (e.g. network 8 – green dotted line), but worsen the outcome in others (e.g. network 3 – blue solid line). These plots indicate that both the random initialization of the network and the different stimulation signals during the CR-on period impact on the final outcome at the end of the CR-off period in a complex manner.

### Impact of RVS CR Stimulation Intensity and Frequency on Acute Effects

Next, we investigate how stimulation intensity and stimulation frequency impact on the mean synaptic weight and synchronization at the end of the RVS CR-on period. Fig 3A shows the median of the mean synaptic weight *C*_*av*_, and Fig 3B of the order parameter *R*_*av*_(averaged over the last 100 · *T*_*s*_) as a function of stimulation intensity *K* and stimulation frequency f_stim_. The color bars show the median values which were calculated from 11 different random initial network configurations. Overall, at the end of the RVS CR-on period we observe a weak excitatory coupling. In other words, CR stimulation shifts the networks’ couplings towards more inhibition, the inhibitory couplings get stronger, and desynchronized states emerge for most of the (*K*, *r*_0_) pairs, except for the two columns at f_stim_ = 25%f_0_(*T*_*s*_ = 64 ms) and f_stim_ = 145%f_0_(*T*_*s*_ = 11 ms). For the former frequency, CR stimulation fails to weaken both the inter-neural connectivity and synchrony, whereas for the latter frequency CR down-regulates synaptic connectivity, but elevated levels of synchrony persist. Figs 3C and 3D show their Inter-Quartile-Range (IQR) respectively, which gives a measure of the data dispersion around these median values. All IQR values being close to zero indicate that the middle 50% of the distribution are very close to the median value.

**Fig 3.**
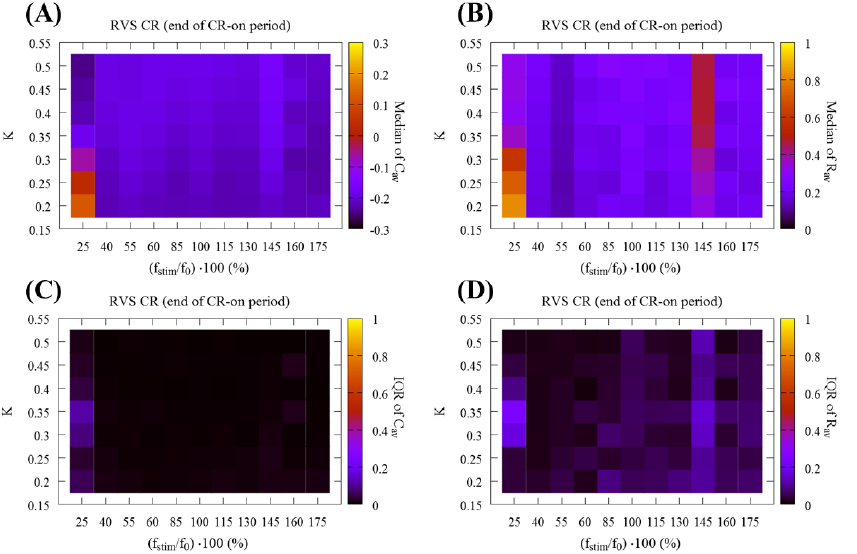
Global overview of the synaptic connectivity and synchronization at the end of the CR-on period using RVS CR stimulation. (A) Median of mean synaptic weight *C*_*av*_ and (B) median of the order parameter *R*_*av*_ at the end of the CR-on period as a function of stimulation intensity *K* and stimulation frequency ratio *r*_0_ = (f_stim_/f_0_) · 100. Color-bars show the median values which were calculated from 11 different random initial network configurations. Panels (C) and (D) show the corresponding IQR, which gives a measure of the dispersion around these median values. All IQR values being close to zero indicate that the middle 50% of the distribution are very close to the median value.

### Impact of RVS CR Stimulation Intensity and Frequency on Sustained After-Effects

Fig 4 presents a global overview of the long-lasting impact of CR at the end of the CR-off period. Fig 4A shows the median of the mean synaptic weight *C*_*av*_, and Fig 4B the median of the order parameter *R*_*av*_. Figs 4C and 4D display the corresponding IQRs, showing that the dispersion around the median of the *C*_*av*_ results is very small in large parts of the parameter plane. In contrast, small IQRs are found only for small *R*_*av*_, in regions with strong desynchronization. Figs 4A and 4B display two main bands in the (*K*, *r*_0_) - parameter space associated with small dispersion: The first band is aligned along the horizontal axis, for weak stimulation intensities (i.e. *K* = 0.20 and *K* = 0.25) and stimulation frequencies greater than 40% of the standard f_0_ corresponding to a stimulation period of *T*_0_ = 16 ms. The second band runs along the vertical stimulation intensity *K* axis, and for relatively high frequencies, i.e. for f_stim_ = 160%f_0_ (*T*_*s*_ = 10 ms) and f_stim_ = 175%f_0_ (*T*_*s*_ = 9 ms) which correspond to ∼ 155% and ∼ 140% of the firing rate of the synchronized neurons, respectively. For these (bottom-horizontal and right-hand-side-vertical bands) the dispersion around the median values is quite small for both *C*_*av*_ and *R*_*av*_(Figs 4C and 4D). In addition, the vertical stripe at the reference frequency value f_0_ (”100%”, *T*_*s*_ = 16 ms), studied in [78], but with a *t* = 64 s CR-on period, is also associated with robust long-lasting anti-kindling and desynchronization for all CR stimulation intensity values *K*. Another region with similar characteristics lies at the center of Figs 4A and 4B for intermediate stimulation intensity and frequency values.

**Fig 4.**
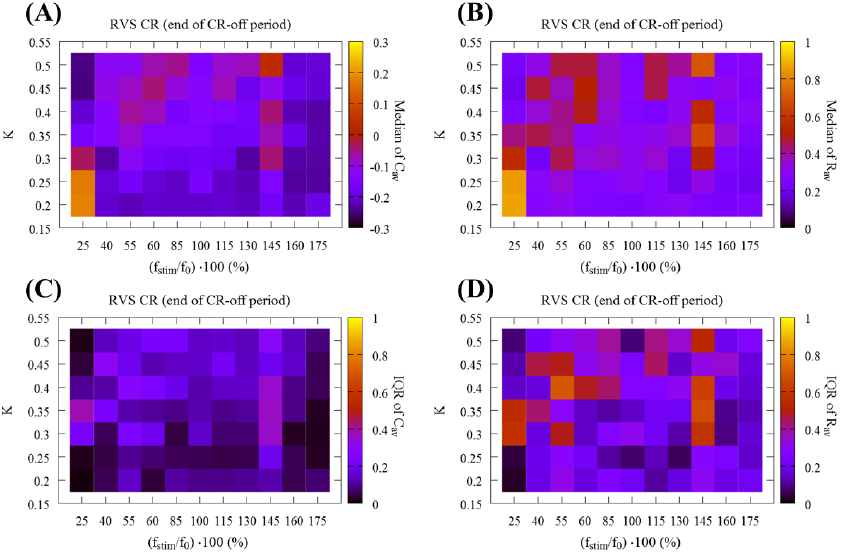
Global overview of the mean synaptic weight and synchronization at the end of the CR-off period using RVS CR stimulation. (A) Median of the mean synaptic weight *C*_*av*_, (B) median of the order parameter *R*_*av*_(11 different random initial network configurations and 11 different RVS CR random signals). Long-lasting anti-kindling is achieved in all dark regions as indicated by the corresponding color-bars. Panels (C) and (D) show the dispersion around these median values by plotting their IQR respectively. All IQR values being close to zero indicate that the middle 50% of the distribution are very close to the median value.

At a first glance, among those two bands in Figs 4A and 4B, where dark color dominates suggesting long-lasting anti-kindling after cessation of CR stimulation, the horizontal band seems especially intriguing. Along the lines of our model analysis, the horizontal band corresponds to pronounced desynchronizing outcome at favorably weak CR stimulation intensities within a range of stimulation frequencies. However, we have to keep in mind that the discrete grid is not very dense. Hence, in order to investigate whether this conclusion is justified, we calculated *C*_*av*_ and *R*_*av*_ for all the integer period *T*_*s*_ values for *K* = 0.20, ranging from f_stim_ = 175%f_0_ (*T*_*s*_ = 10 ms) to f_stim_ = 40%f_0_ (*T*_*s*_ = 40 ms). Fig 5 shows this fine-grained analysis. The boxplot for *C*_*av*_ is shown in Fig 5A, and for the *R*_*av*_ in Fig 5B. Note, in this figure the horizontal axis shows the CR stimulation period instead of the frequency. And it is sorted from larger to smaller values for an easier comparison between the two representations. The red and green dots indicate the reference stimulation period *T*_0_ = 16 ms and intrinsic firing rate period *T*_int_ = 14 ms respectively. For *T*_*s*_∈ [9, …, 24 ms] we observe robust anti-kindling effects. In contrast, for *T*_*s*_∈ [25, …, 28] many networks tend to be in a synchronized state, while for *T*_*s*_∈ [29, …, 38] the anti-kindling is found to be robust again, before finally reaching the largest *T*_*s*_ value where the CR stimulation signals are not effective at all. In summary, at weak stimulation intensities favorable stimulation outcomes are achieved within wide ranges of the stimulation frequency. For further analyses of stimulation induced effects observed in particular ranges of the stimulation intensity/frequency parameter plane, we refer to the *Supporting Information*. For particular stimulation parameters, similar acute effects, as assessed with macroscopic quantities *R*_*av*_ and *C*_*av*_, may lead to qualitatively different results. Neither prominent features of the connectivity matrix nor the dynamical states of the individually stimulated subpopulations at the end of the CR-on period enabled us to predict the long-term outcome (see *Supporting Information*). Furthermore, this analysis revealed that CR may be effective without causing side-effects that are time-locked to the individual stimuli (see *Supporting Information*).

**Fig 5.**
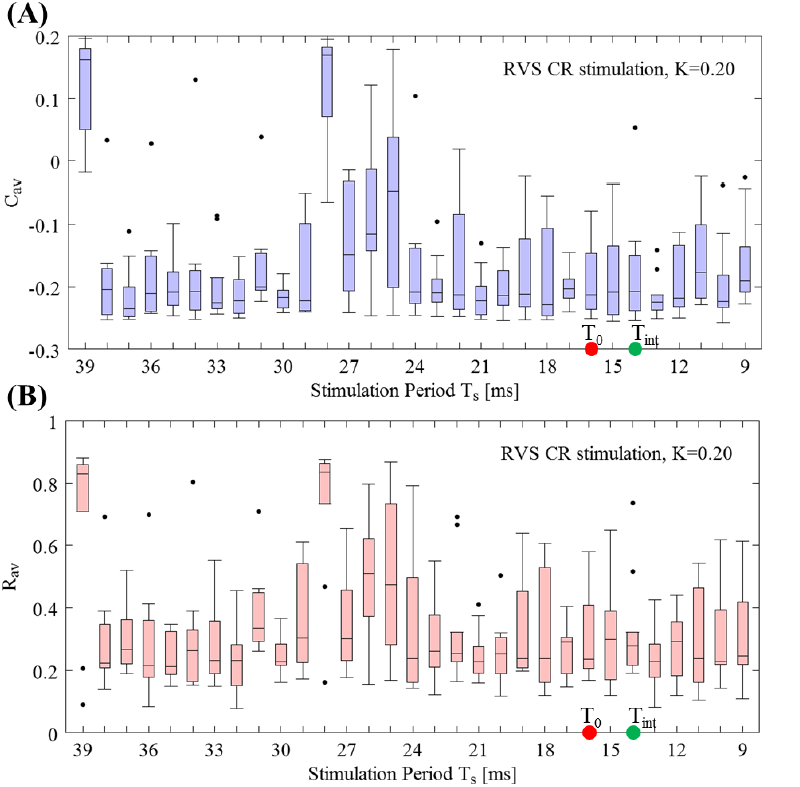
Fine-grained *T*_*s*_-period grid analysis for RVS CR stimulation at intensity *K* = 0. 20. (A) Boxplots of *C*_*av*_(mean synaptic weight) and (B) *R*_*av*_(order parameter) for fixed and weak stimulation intensity *K* = 0.20 for a finer sample on the *T*_*s*_ integer value interval at the end of the CR-off period. The red and green dots indicate the reference stimulation period *T*_0_ = 16 ms and intrinsic firing rate period *T*_int_ = 14 ms, respectively.

### Impact of SVS CR Stimulation Intensity and Frequency on Stimulation Effects

Next, we address the robustness of the long-lasting anti-kindling achieved by SVS CR stimulation in the (*K*, *r*_0_) -parameter plane. We use SVS-100 CR stimulation, where the random switching occurs after 100 repetitions of the CR sequence (for motivation see [78]). In Fig 6 we show the total outcome of *C*_*av*_ and *R*_*av*_, obtained by delivering SVS-100 CR to the same 11 initial networks as in Figs 3 and 4 and varying the CR stimulation frequency and intensity. Let us compare these results with the results for RVS CR (Figs 3A and 3B). Regarding the medians of the *C*_*av*_, both RVS and SVS CRs (Fig 3A vs Fig 6A) overall the parameter dependence outcomes are similar, where the outcome plots of SVS CR (Fig 6) contain more vertical stripes, associated with greater outcome variability. Let us consider some of the differences between RVS CR and SVS CR: For low intensity (*K* = 0.20) and high frequencies f_stim_ = 175%f_0_, 160%f_0_ (corresponding to *T*_*s*_ = 9 ms, 10 ms respectively) SVS-100 does neither cause pronounced acute desynchronizing effects nor sustained long-lasting effects. For low CR frequency 25%f_0_ (corresponding to *T*_*s*_ = 64 ms, leftmost column) it requires even stronger intensities to induce an anti-kindling compared to RVS (Fig 3A). Regarding the median of *R*_*av*_(Fig 3B vs Fig 6B) for almost all (*K*, *r*_0_) -parameters the networks are shifted to a desynchronized state at the end of the CR-on period, with only a few exceptions, in particular (*K*, *r*_0_) = (0.20, 175%f_0_), (0.20, 160%f_0_) and for 25%f_0_. Moreover, for the frequency f_stim_ = 145%f_0_ the SVS CR stimulation achieves more pronounced anti-kindling effects (at the end of the CR-on period) for all intensities *K* compared to the RVS CR stimulation.

**Fig 6.**
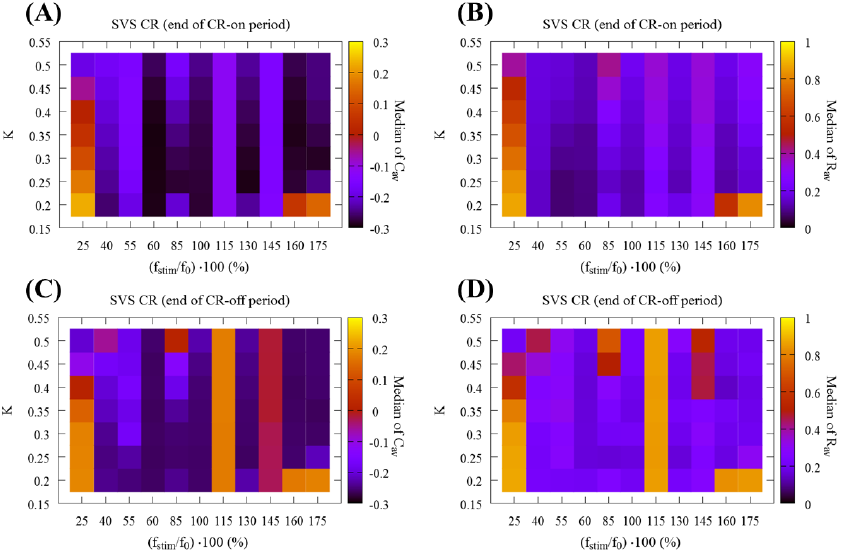
Global overview of the mean synaptic weight and synchronization at the end of the CR-on/off period using SVS CR stimulation. (A) Median of the mean synaptic weight *C*_*av*_ and (B) median of the order parameter *R*_*av*_ for 11 different random initial network configurations at the end of the CR-on period. (C) Median of the mean synaptic weight *C*_*av*_ and (D) median of the order parameter *R*_*av*_ at the end of the CR-off period.

In Figs 6C and 6D we present the outcome for the medians of *C*_*av*_ and *R*_*av*_ at the end of the CR-off period for SVS-100 CR. The main differences compared to RVS CR (Figs 4A and 4B) are the ‘stripes’ at f_stim_ = 115%f_0_ and, in particular, at f_stim_ = 145%f_0_ where SVS-100 neither reduces *C*_*av*_ nor *R*_*av*_ for all *K*-values. Moreover, also for the lowest intensity value *K* = 0.20 and frequencies f_stim_ = 175%f_0_, 160%f_0_, 25%f_0_ no anti-kindling is achieved. However, there is a substantial overlap of the (*K*, f_stim_) -parameter range where both RVS and SVS-100 CR lead to long-lasting anti-kindling, mainly for high frequencies f_stim_ = 175%f_0_, 160%f_0_ for *K* ≥.0 25 as well as for 40%f_0_ ≲ f_stim_≲ 100%f_0_ and a wide range of *K*-values. Interestingly, whenever SVS CR stimulation causes an anti-kindling, the long-term effects on the connectivity are particularly robust, irrespective of different network initializations and parameters (data not shown here).

Let us now investigate a denser *T*_*s*_ period sample for the weakest intensity *K* = 0.20, with the same format as in Fig 5 for RVS CR stimulation. Boxplots of *C*_*av*_(Fig 7A) and *R*_*av*_(Fig 7B) at the end of the CR-off period show that SVS CR stimulation at this weak intensity is overall less efficient in inducing long-lasting anti-kindling effects compared to RVS CR (Fig 5). In particular, there is no distinct range of *T*_*s*_ periods where SVS CR causes a pronounced anti-kindling. However, for a few values of *T*_*s*_ for the long-term outcome for SVS is stronger than for RVS, e.g. for *T*_*s*_ = 15 ms, 16 ms.

**Fig 7.**
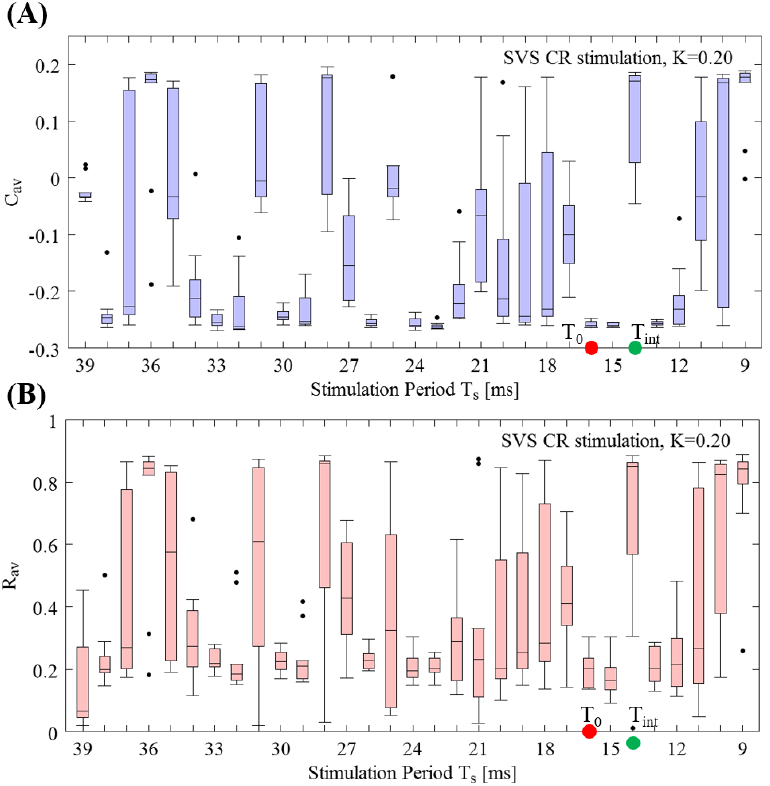
Fine-grained *T*_*s*_-period grid analysis for SVS CR stimulation at intensity *K* = 0. 20. (A) Boxplots of *C*_*av*_(mean synaptic weight) and (B) *R*_*av*_(order parameter) for fixed and weak stimulation intensity *K* = 0.20 for a finer sample on the *T*_*s*_ integer value interval at the end of the CR-off period. The red and green dots indicate the reference stimulation period *T*_0_ = 16 ms and intrinsic firing rate period *T*_int_ = 14 ms, respectively. Format as in Fig 5.

## Discussion

By systematically varying the CR stimulation frequency and intensity and comparing the stimulation outcome of the two different CR protocols, RVS and SVS CR stimulation, RVS CR proved to be more robust with respect to variations of the stimulation frequency. However, in accordance with a previous computational study, restricted to a fixed value of the stimulation frequency [78], SVS CR stimulation can induce stronger anti-kindling effects. In our study, we obtained particularly parameter ranges related to particularly favorable stimulation outcome. If no closed loop adaptation for the stimulation frequency is available, RVS CR stimulation at weak intensities and with stimulation frequencies in the range of the neuronal firing rates enables to effectively and robustly achieve an anti-kindling.

To our knowledge, in our study in a plastic network the CR stimulation frequency and intensity were systematically varied for the first time to investigate the impact on acute and long-lasting stimulation outcome. Remarkably, pronounced acute desynchronization (as measured by means of the standard order parameter from Eq (11) [76, 103]) does not necessarily lead to long-lasting desynchronization. On the one hand this finding might inspire future computational and pre-clinical studies aiming at specifically designing stimulation protocols for long-lasting (as opposed to acute) desynchronization. On the other hand, this finding is significant for the development of clinical calibration procedures for CR stimulation, see [114].

In a previous study in networks without STDP Lysyansky and coworkers [75] considered *m*: *n* ON-OFF CR stimulation with real rather than integer *m* and *n* and varied *m* and *n* systematically. For non-integer *m* incomplete CR stimulation cycles are delivered, intersected by incomplete pause cycles caused by non-integer *n*. This type of CR stimulation has not yet been used in pre-clinical or clinical studies and is somewhat remote to the initial CR concept that builds on the periodicity of both neuronal firing and stimulus patterns [58, 59].

For the majority of the CR stimulation parameters used in this work, no drastic change was observed in the firing rates (data not shown). The only exception was observed for very low stimulation frequency combined with comparably high intensities (S2 Fig). Especially for the most relevant cases (weak to intermediate intensities and frequencies around the reference stimulation frequency) the firing rate of the neuron ensemble remains almost unchanged when compared with initial intrinsic firing rates before CR delivery (about up to ±**3**% variation of the initial intrinsic firing rate). In this study, we focused on a network of spiking Hodgkin-Huxley neurons with STDP. Compared to STDP-free networks used before [75], this is a step towards more complex and, in particular, plastic neural networks. Future studies should address yet more complex neural networks equipped with STDP to study parameter regions and stimulation protocols that are reasonably stable in different neural network models. In principle, we have to be careful about extrapolating findings obtained in one type of neural network model to network models of higher complexity. For instance, non-linear delayed feedback stimulation was introduced in globally coupled networks of limit cycle oscillators and phase oscillators [106]. It turned out to robustly cause desynchronization, nearly irrespective of the selected valued of the delay [107]. In contrast, linear delayed feedback [108] was shown to induce desynchronization only for a rather small subset of parameter pairs of delay and intensity, favoring delays close to half the intrinsic oscillation period and weak to moderate intensities [107, 108].

However, in a more complex, microscopic neuronal network model consisting of a population of STN and a population of external globus pallidus (GPe) neurons [99] the parameter dependence for nonlinear delayed feedback was qualitatively different [109]. The parameter ranges of delay and intensity values associated with desynchronization were still greater for nonlinear delayed feedback as opposed to linear delayed feedback. However, in this microscopic STN-GPe network model nonlinear delayed feedback had to be properly calibrated and, in particular, the delay had to be adjusted to the intrinsic period of the neuronal oscillations, to enable desynchronization [109]. Note, the microscopic STN-GPe network model did not contain STDP [99]. Incorporating STDP to a neuronal network model substantially adds to the model’s complexity (see e.g. [28]) and might, hence, further impact on the dependence of the stimulation outcome on key parameters of both linear and nonlinear delayed feedback. Ultimately, we strive for using several neural networks with STDP as testbed for generating computationally based predictions and recommendations for favorable stimulus parameters and dosage protocols. However, different models may display similar or even identical spontaneous (i.e. stimulation-free) dynamics, but may have very different stimulus response properties (see e.g. [56]).

Accordingly, we cannot expect a stimulation technique to be generically effective, irrespective of the neural network model used. Nevertheless, stepwise adding further physiologically and anatomically relevant features to the neural network models employed may help to generate specific predictions and, ultimately, to further improve stimulation protocols and dosage regimes. In that sense, the finding that RVS CR stimulation at weak to moderate intensities and stimulation frequencies adapted to the neurons’ intrinsic firing rates causes a desynchronization in neural network models without STDP [75] and with STDP as shown in this study, is relevant and, in fact, in accordance with pre-clinical findings [65, 68]. Furthermore, the fact that SVS CR stimulation might even be more effective, but requires more careful parameter adaptation may guide future development of calibration techniques as put forward in a forthcoming paper study [Manos T, Zeitler M, Tass PA., Improving long lasting desynchronization effects via coordinated reset stimulation mild frequency modulation, PLOS Computational Biology (submitted simultaneously) 2017].

In neural networks with STDP post-stimulation transients may be complex. For instance, for stimulation dosages just reaching the level required for an anti-kindling, a rebound of excessive synchrony may occur immediately following cessation of CR stimulation, while later on a full-blown, sustained desynchronization emerges [37, 61, 110]. This rebound selectively relates to synchrony, rather than synaptic connectivity. This phenomenon occurs when after CR delivery the neuronal population just reaches the basin of attraction of a favorable attractor. Upon entering the basin of attraction, the synaptic connectivity is still super-critical, so that synchrony emerges in the absence of stimulation. As the neuronal network relaxes towards the favorable attractor, the initially up-regulated synaptic connectivity fades away until, finally, the synaptic connectivity remains below a critical threshold, hence, preventing the population from getting synchronized [37, 61, 110]. However, it remains to be shown to which extent the rebound of synchrony phenomenon might be a generic after-effect occurring for just about sufficient CR dosage or simply an epiphenomenon specific to the computational model [111] used in those studies [37, 61, 110], comprising networks of Morris-Lecar spike generators [112] transformed to burst mode by a slowly varying current [112, 113].

We here demonstrated that over a wide range of stimulation parameters favorable acute effects do not automatically lead to favorable long-lasting, sustained after-effects. This is in agreement with a computational study in the same model, but performed in only a restricted parameter range [78], as well as with an EEG experiment performed in tinnitus patients [114]. To characterize stimulation induced effects, we here used the average synaptic weight [Eq (10)] and the average amount of neuronal synchrony [Eq (11)]. These macroscopic quantities enabled us to effectively investigate the impact of variations of stimulation parameters on the stimulation outcome. However, in *Supporting Information* section, we have shown that pronounced differences of the average synaptic weight do not necessarily lead to pronounced differences of the average amount of synchrony (S1 Fig). Another example in this context is the combination of weak average synaptic connectivity (Fig 3A) combined with increased levels of average neuronal synchrony (Fig 3B) at the end of the CR-on period. To further study the relationship between connectivity pattern and synchronization processes, macroscopic quantities may not be sufficient to grasp all relevant details of the connectivity matrix and the dynamical features of the resulting synchronization processes.

Though not in the intended scope of our present study, the actual mechanism of action of CR stimulation deserves attention in future studies. In neural and oscillator networks without STDP, CR stimulation disrupts synchrony by causing phase resets of different subpopulations at different times [58, 59, 75]. The phase reset of a single subpopulation is time-locked to the stimulus affected that particular subpopulation [58, 59, 75]. However, in the present study we observed that CR stimulation may not just reorder the neurons’ phases. Rather, for particular stimulation parameters it may even cause a significant decrease of the neuronal firing rates, intriguingly associated with a particularly pronounced anti-kindling (S2I Fig). Furthermore, in contradiction to the results obtained in networks without STDP [58, 59, 75], CR stimulation may cause a full-blown anti-kindling without any phase resets of the subpopulations time locked to the corresponding stimuli (S4 Fig). This is relevant for two reasons: (i) Since effective CR stimulation does not require phase resets time-locked to the individual stimuli, further computational studies should elucidate whether it makes sense to calibrate CR stimuli for pre-clinical and clinical applications by selecting stimulus parameters that favorably achieve phase resets. Corresponding results might be relevant for the design of calibration procedures and, in addition, challenge existing patents that are based on selecting parameters that optimally achieve phase resets of the stimuli delivered to the individual sub-populations (e.g. [115]). (ii) By the same token, our results do not only challenge current hypotheses on the mechanism of CR stimulation, but also fundamental patents in the field of invasive (e.g. [116]) as well as non-invasive (e.g. [117]) CR stimulation. Accordingly, future computational studies should focus on the mechanism of action of CR stimulation in networks with STDP in order to actually understand and possibly improve anti-kindling protocols.

Our goal is to accomplish an anti-kindling in a way as robust as possible, complying with clinically motivated constraints. For instance, striving for anti-kindling induced at minimal stimulation intensities led to the computational development of spaced CR stimulation [118] and two-stage CR stimulation with weak onset intensity [79]. The motivation behind these developments was to avoid side effects by substantially reducing stimulation intensities [79, 118]. Another direction is to accomplish anti-kindling at moderate stimulation duration as computationally studied in this paper. This may be favorable from a clinical standpoint since it might help to reduce the occurrence of side effects as well as the requirement of the treatment on patients for their compliance, e.g., in terms of actually using non-invasive therapeutic devices. In this context, it might turn out to be beneficial that RVS CR stimulation causes sustained after-effects over a wide range of stimulation frequencies even at weak intensity (Fig 4). Accordingly, RVS CR stimulation might provide an appropriate stimulation protocol, in particular, if applied in an open-loop manner, without the ability to calibrate the stimulation parameters, especially the stimulation frequency by adapting it to the dominant peaks in the frequency spectrum of electrophysiological signals such as local field potentials or EEG signals. However, in a forthcoming computational study [Manos T, Zeitler M, Tass PA., Improving long lasting desynchronization effects via coordinated reset stimulation mild frequency modulation, PLOS Computational Biology (submitted simultaneously) 2017] we will use comparably simple closed-loop control modes to significantly improve the robustness of both RVS and SVS CR stimulation and, in particular, exploit the anti-kindling potential of SVS CR stimulation.

## Author Contributions

Conceived and designed the experiments: TM MZ PAT. Performed the experiments: TM. Analyzed the data: TM MZ PAT. Contributed reagents/materials/analysis tools: TM MZ. Wrote the paper: TM MZ PAT.

## Supporting Information for

### ”How stimulation frequency and intensity impact on the long-lasting effects of coordinated reset stimulation”

Thanos Manos, Magteld Zeitler and Peter A. Tass

### Relationship between Average Synaptic Weight and Average Synchrony

In S1 Fig we focus on the three prominent vertical stripes connected with favorable stimulation outcome (indicated by the white arrows in the insets of S1 Fig), where the median values for both mean synaptic weight and order parameter are small, indicating weak excitatory connectivity and synchrony. Along all these three stripes CR induces an anti-kindling (see S1A-C Figs) and a long-lasting desynchronization (see S1D-F Figs). For *T*_*s*_=9 ms, 10 ms the impact of CR is stronger compared to *T*_*s*_ = 16 ms and leads to weaker synaptic connectivity and further reduction of synchrony. However, the latter difference does not translate to the amount of synchrony. In fact, for some *K* values the period *T*_*s*_ = 16 ms we observe a more desynchronized state than *T*_*s*_ = 9 ms, 10 ms (S1D-F Figs). Put otherwise, pronounced differences of the average synaptic weight need not be reflected by pronounced differences of the average amount of synchrony.

**S1 Fig.**
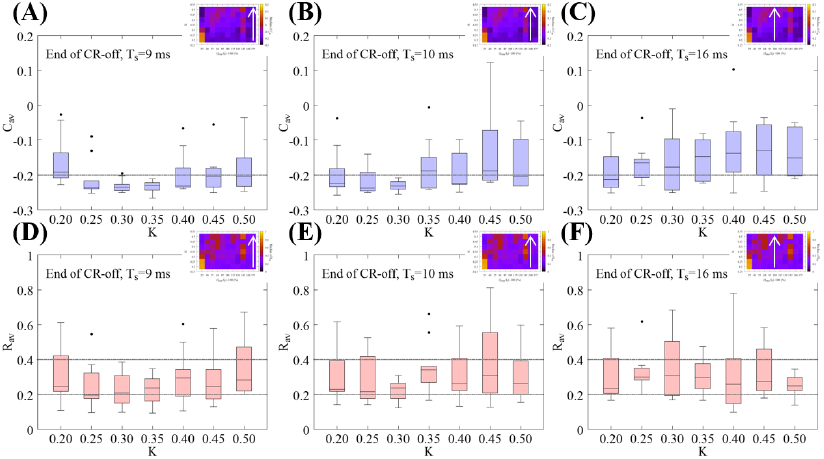
Boxplots for the mean synaptic weight *C* _*av*_ and the order parameter *R*_*av*_ for stimulation periods with long-lasting anti-kindling at all RVS CR stimulation intensities *K*. (A) *C*_*av*_ and (D) *R*_*av*_ for *T*_*s*_ = 9 ms. (B) *C*_*av*_ and (E) *R*_*av*_ for *T*_*s*_ = 10 ms. (C) *C*_*av*_ and (F) *R*_*av*_ for *T*_*s*_ = 16 ms. These three period values are indicated by the white arrows in the inset *C*_*av*_ and *R*_*av*_ general median overview color-plots. The dotted horizontal line(s) (one for the *C*_*av*_ and two for the *R*_*av*_ boxplots) are visual cues to facilitate comparison between different panels.

### Intensity Dependent Stimulation Effects

At the stimulation frequency f_stim_ = 145%f_0_(*T*_*s*_ = 11 ms) for larger stimulation intensities no pronounced desynchronization is achieved, and acute effects (Fig 3) as well as sustained aftereffects are poor (Figs 4A and 4B). A qualitatively different phenomenon occurs at the lowest stimulation frequency f_stim_ = 25%f_0_ (*T*_*s*_ = 64 ms): The anti-kindling effects get more pronounced with increasing stimulation intensity. S2 Fig presents a detailed analysis for different stimulation intensities *K*, where S2A and S2B Figs depict boxplots of *C*_*av*_ and *R*_*av*_ for the different *K* values. A clear inverse *sigmoidal*-like decay is observed in both figures indicating a monotonic-like gradual tendency to more effective long-lasting anti-kindling as the CR intensity increases for this particular stimulation period. The time evolution of *C*_*av*_ for each different network is shown in S2C-F Figs for *K* = 0.20, 030, 0.40, 0.50. The most striking result is presented in S2F Fig with *K* = 0.50 where we observe an optimal impact of CR stimulation, i.e. all networks/signals reach a rather low value and maintain this low mean synaptic weight value until the end of the CR-off period.

**S2 Fig.**
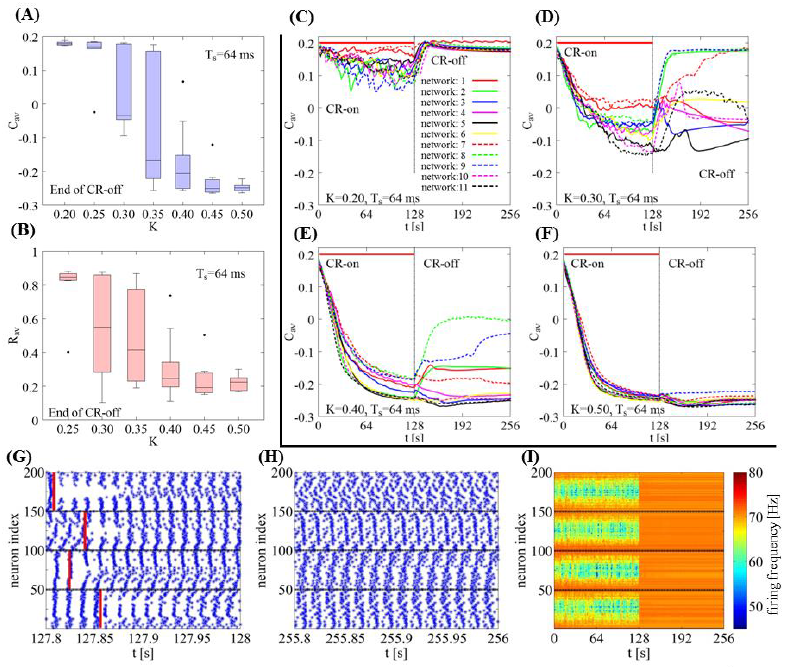
Detailed analysis of the *T*_*s*_ = 64 ms period for different stimulation intensity values *K*. (A) Boxplots of *C*_*av*_ for the different *K* values and (B) boxplots for *R*_*av*_. (C)- (F) Time evolution of *C*_*av*_ for *K* = 0.20, 030, 0.40, 0.50 and 11 different initial networks and signals. (G) Raster plot at the end of the CR-on period for *K* = 0.50 and network 1. (H) Raster plot at the end of the CR-off period. Blue points indicate the spiking times (horizontal axis) of each neuron (vertical axis), the red vertical lines point out the onset of each CR stimulation at the center of these sub-intervals (neurons *i* = 25, 75, 125, 175). (I) Firing frequencies of the individual neurons for *K* = 0.50 (network 1). The horizontal black lines (G, H and I) are visual clues to distinguish between the four stimulated neuronal subpopulations.

S2G and S2H Figs show the raster plots at different time windows (each of 200 ms width) for the *K* = 0.50 (network 1) case of S2F Fig, at the end of the CR-on and at the end of the CR-off period respectively. The horizontal black lines are visual cues, distinguishing the four separately stimulated groups of neurons. CR stimulation is administrated at neurons *i* =25, 75,125,175. S2I Fig depicts the impact of the CR stimulation, again for *K* = 0.50 (network 1), on the firing frequencies (color bar) of the network, calculated in time windows of length 100 · *T*_*s*_. Let us recall (see *Simulation description* section) that the intrinsic fire rate before the CR is ~71 Hz. Hence, the CR stimulation, in this particular effective case, causes a decrease in firing ratet to ?60Hz (greenish range).

### Predictability of Long-Term Outcome and Mechanism of CR Stimulation on the Presence of STDP

This section serves to illustrate two findings: (i) Macroscopically similar states at the end of the CR-on period may lead to qualitatively different long-term dynamics. (ii) Neither prominent features of the synaptic connectivity matrix at the end of the CR-on period nor the dynamics of the selectively stimulated subpopulations at the end of the CR-on period enabled us to predict the long-term outcome.

To investigate the impact of different initial network conditions and different realizations of RVS CR sequences on the long-lasting sustained effects, we consider one parameter pair (*K*, *T*_*s*_) and two different network initializations. For both networks 1 and 2 the mean synaptic weight *C*_*av*_ reaches similar, low values by the end of the CR-on period (S3A Fig) with similar connectivity matrices (S3B and S3D Figs). Nevertheless, in both cases the long-term outcome is qualitatively different concerning both the mean synaptic weight *C*_*av*_(S3A Fig) and the corresponding connectivity matrices (S3C and S3E Figs).

**S3 Fig.**
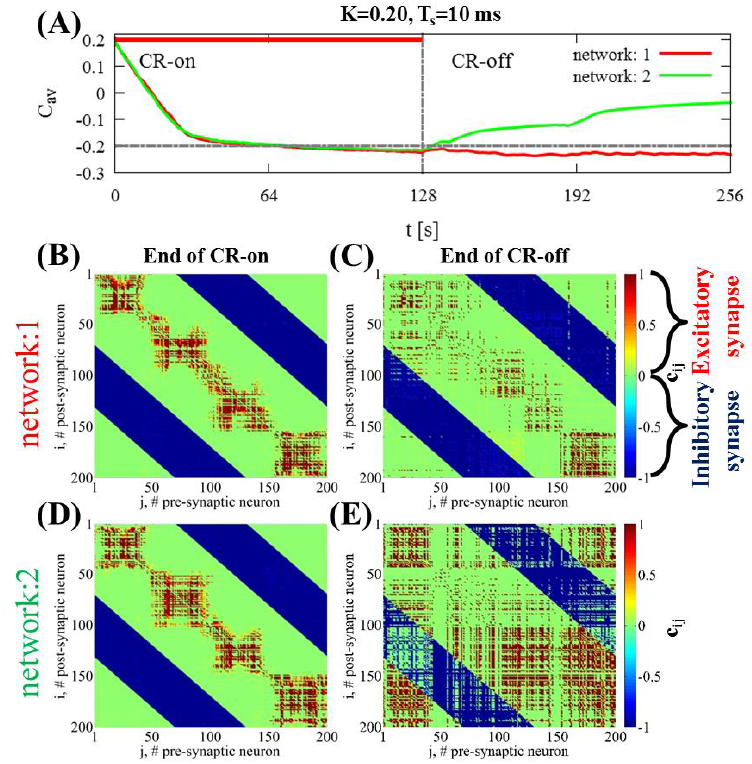
Connectivity at the end of the RVS CR-on period and related long-term outcome. (A) The time evolution of the mean synaptic weight *C*_*av*_ and the corresponding connectivity matrices at the end of the CR-on period [panels (B) and(D)] and CR-off period [panels (C) and (E)] for two different network initializations and random RVS CR sequences: for network 1 [red solid line in (A) and connectivity matrices (B), (C)] and network 2 [green solid line in (A) and connectivity matrices (D), (E)] for *K* = 0.20, *T*_*s*_ = 10 ms.

In addition, we consider the amount of synchrony of the entire neural network and its subpopulations as well as the corresponding raster plots for the same set of parameters as in S3 Fig. S4A Fig shows the moving average of the order parameter < *R* > (in a running window of 400 · *T*_*s*_ length) for the whole neuron ensemble. We use moving averages because of the presence of strong fluctuations of *R*. At *t* = 0 s both networks are strongly synchronized (*R* ≈ 0.85) and rapidly undergo a CR-induced desynchronization. After the cessation of CR stimulation, both networks maintain a similar degree of global desynchronization for up to *t* ∼190 s, when only one of them starts getting more synchronized (green solid line, network 2). CR is delivered at four stimulation sites. To specifically analyze the impact of CR on the subpopulations in the vicinity of the stimulation sites, we calculated the time average of the order parameter for each subpopulation, too.

**S4 Fig.**
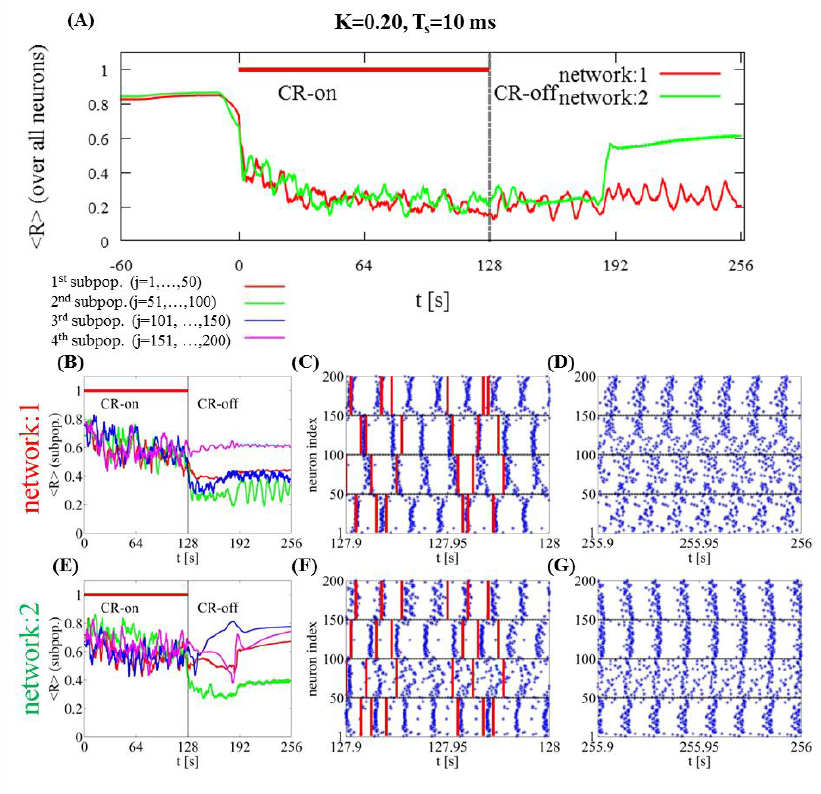
Impact of different networks/signals on the predictability of the long-lasting desynchronization. (A) Moving average of the order parameter < *R* > (averaged over a sliding window of 20 ms width and *K* = 0.20, *T*_*s*_ = 10 ms) for the whole network and for the four subpopulations of networks 1 (B) [belonging to network 1 of (A)] and 2 (E) [network 2 of (A)]. The definition of the four subpopulations reflects the equidistant delivery of CR stimulation and the corresponding order parameters are coded by different color (see inset). In the raster plots at the end of the CR-on period (C), (F) and at the end of the CR-off period (D), (G), blue points indicate spiking times (horizontal axis) of each neuron (vertical axis), red lines illustrate the onset of each CR stimulus at the center of that neuronal subpopulation. Horizontal black lines delineate the allocation of the four neuronal sub populations.

In the second and third row of S4 Fig we show the time averaged order parameter < *R* > for each subpopulation. The allocation to subpopulations is given by the stimulation setup, as CR stimuli are equidistantly delivered to four different sites. The subgroups’ order parameters are indicated by different colors (S4B and S4E Figs) for network 1 (red solid line in S4A Fig) and 2 (green solid line in S4A Fig) respectively. The amount of synchrony within the four subpopulations is similar during stimulation, but quite different post-stim. In network 1 three of the subpopulations remain in a desynchronized state post-stim (S4B Fig) and the corresponding neurons do not fire coincidently (S4D Fig). In contrast, three of the subpopulations of network 2 resynchronize (S4E Fig), as reflected by the coincident spiking of their neurons (S4G Fig).

This example is intended to illustrate the multitude of dynamic states evolving from comparably similar states at stim offsets. These different states differ with respect to extent of synchrony within subpopulations, their mutual phase relationship and time courses of these quantities. We found several parameter pairs (*K*, *T*_*s*_) with similar acute, but qualitatively sustained long-lasting effects. For these different parameter pairs, we performed similar comparisons as in the example above. Macroscopic quantities like *C*_*av*_ and < *R* > of the entire network or of the subpopulations together with the subpopulations’ mutual phase relationships did not enable us to find markers predictive of specific long-term outcome in cases with a plurality of CR-off responses originating from similar end of CR-on states.

Intriguingly, RVS CR stimulation does not induce phase resets of the individual subpopulations that are time-locked to the corresponding stimuli (S4C and S4F Figs). Rather, the time differences between the stimulus onsets (red bars in S4C and S4F Figs) and the neurons’ spikes (blue dots in Figs S4C and S4F) do not cluster around a preferred value. This finding was repeatedly observed in a portion of parameter pairs (*K*, *r*_0_).

